# The heat is on: behavioural, physiological and reproductive evidence of heat stress in breeding king penguins

**DOI:** 10.1101/2024.09.09.611977

**Authors:** Aude Noiret, Agnès Lewden, Camille Lemonnier, Céline Bocquet, Marine Montblanc, Fabrice Bertile, Marine Hoareau, Elsa Marçon, Jean-Patrice Robin, Pierre Bize, Vincent A Viblanc, Antoine Stier

**Affiliations:** Université de Strasbourg, CNRS, IPHC UMR 7178, F-67000 Strasbourg, France; Laboratoire d’Ecologie des Hydrosystèmes Naturels et Anthropisés, CNRS UMR 5023 Université Claude Bernard Lyon 1, Université de Lyon, Lyon, France; Université de Brest - UMR 6539 CNRS/UBO/IRD/Ifremer, Laboratoire des sciences de l’environnement marin - IUEM - Rue Dumont D’Urville - 29280 - Plouzané; Swiss Ornithological Institute, CH-6204 Sempach, Switzerland; Department of Biology, University of Turku, Turku, Finland

**Keywords:** thermoregulation, temperature, seabird, climate change, behaviour

## Abstract

Polar and sub-polar animals evolved to thrive in cold climates and may thus be particularly sensitive to rising temperatures associated with climate change. Penguins may be especially vulnerable, due to their dual habitat, alternating between foraging in cold waters and breeding/moulting on an increasingly warm land. Here, we characterized heat stress occurrence in breeding king penguins through behavioural observations (*e.g.* panting occurrence) and body temperature measurements. We observed that behavioural signs of heat stress are frequent in king penguins breeding in the sub-Antarctic region (*>* 20% of observations at mid-day), and that subcutaneous temperatures increase under high heat load, especially in penguins observed panting. Subcutaneous and core body temperatures were moderately correlated and both increased with heat load. Yet, their responses were not parallel since core body temperature is markedly less sensitive to heat load than subcutaneous temperature. Air temperature alone was a poor predictor of heat stress occurrence, whereas the combination of high solar radiation, low wind speed and high air temperatures provided the strongest predictive power. Finally, reproductive failures were more likely to occur on warmer days, suggesting that heat stress may have significant sublethal effects on adults that could ultimately affect population dynamics.

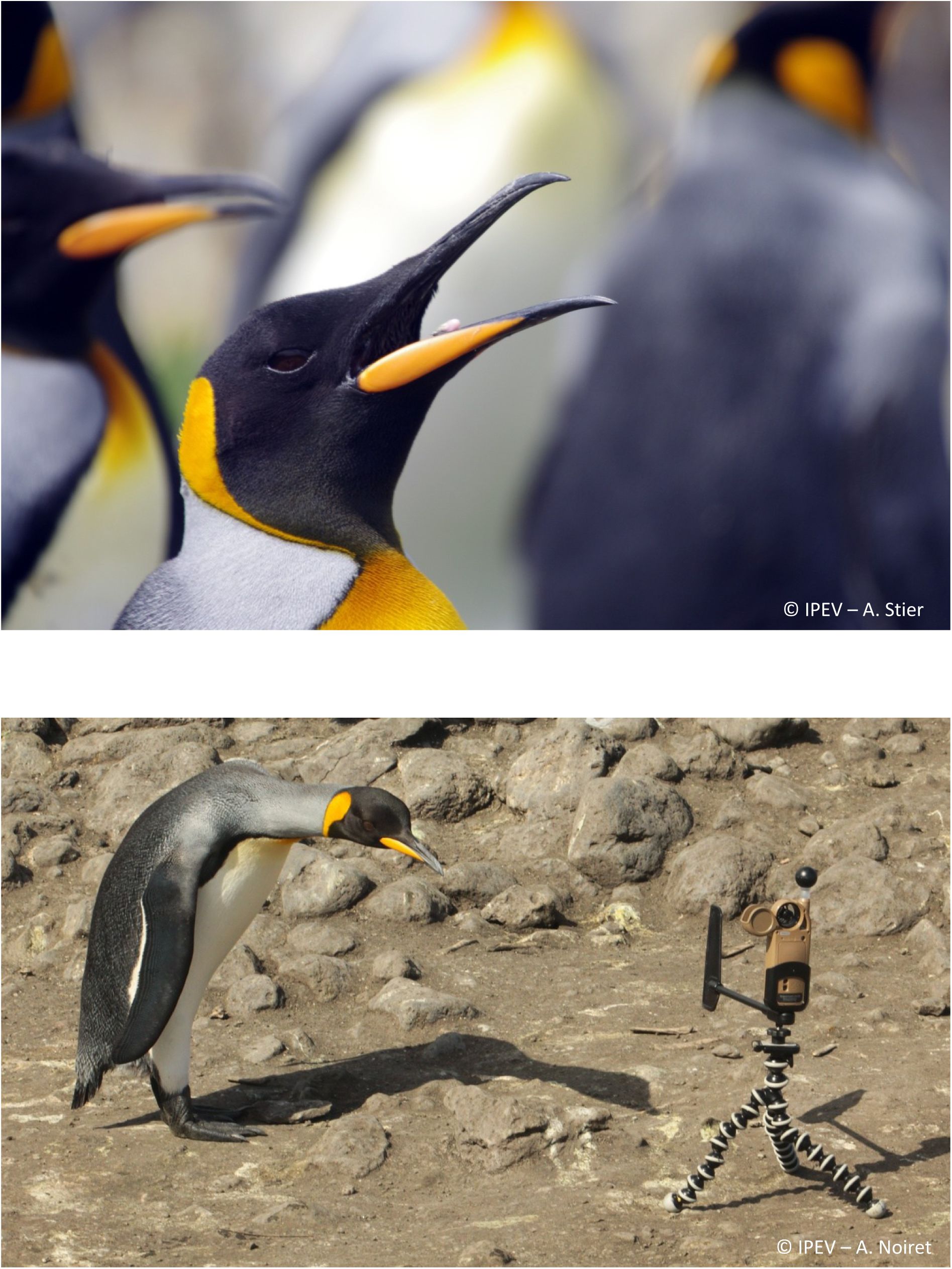

## Introduction

As climate is warming, it is expected to exert direct thermoregulatory pressure on most living organisms (Piatt *et al*. 2024; Stillman 2019). Although heat waves - which are known to cause mass mortality events amongst wild animals (Conradie *et al*. 2020) – are defined on different criteria between studies (Gimeno *et al*. 2025; Robinson 2001; Ton *et al*. 2021), climate can be defined as “hot” as soon as it triggers responses of the organism to maintain or restore body temperature and homeostasis. When the response is energetically costly, involving physiological and behavioural changes, it can be defined as heat stress (Hansen 2009). Indeed in endotherms, when ambient temperatures reach the upper limits of their thermoneutral zone (TNZ) (*i.e.* the range of ambient temperatures in which metabolic rate is minimal) (Romanovsky 2007, 2018), the costs of active thermoregulation or the failure to properly thermoregulate can impair organism functions. This can lead for instance to sublethal effects such as impaired cognitive capacity, metabolism, immune response, reproduction, or growth (Boyles *et al*. 2011; Danner *et al*. 2021; Kim *et al*. 2025; Pipoly *et al*. 2013; Rostagno 2020), but also to lethal effects in extreme situations (Holt & Boersma 2022; McKechnie *et al*. 2021; McKechnie & Wolf 2010). However, as the predicted impact of global warming on animal populations are based primarily on air temperatures (T_air_), the increase in heat load faced by animals may be underestimated, as it depends on multiple climatic parameters, including T_air_, but also solar radiation (SR), wind speed (WS) and humidity (Mitchell 2024). Indeed, heat is gained from solar radiation absorption by the animal’s body or by the surrounding surfaces that conduct additional heat, wind favours heat loss by convection, and humidity increases heat loss at low T_air_ but impedes heat dissipation by evaporation at high T_air_. Studying the effects of these variables is even more relevant since low cloud coverage (reflecting back solar radiation) and wind speed show some regional and seasonal trends of decreasing (Andres-Martin *et al*. 2024; Ceppi *et al*. 2026). Hence, when available, integrative climatic data can be used to compute various thermal indexes (Ioannou *et al*. 2022), which may help to refine the determination of the threshold beyond which an animal could be impacted by “hot” conditions, especially in complex natural environments.

Studies on heat stress occurrence in wild animals are mostly conducted in species from arid, tropical or temperate regions, because of obvious extreme ambient temperatures (Bryant 2008; Conradie *et al*. 2020). Yet, cold-adapted species may be at greater risks of struggling under hot conditions because of a narrower, cold-shifted TNZ and thermoregulatory mechanisms specialized for cold environments (Beaman *et al*. 2024; Choy *et al*. 2021; Grémillet *et al*. 2012; O’Connor *et al*. 2021). Even if maximum temperatures in polar regions are seen as less extreme than in other parts of the globe, polar regions are warming at a faster rate than any other place on earth (Clem *et al*. 2020; Stammerjohn & Scambos 2020; You *et al*. 2021). Hence, animals adapted to cold environments may reach a breakpoint in thermoregulation efficacy more rapidly than other species in response to climate anomalies (Beaman *et al*. 2024; Choy *et al*. 2021; O’Connor *et al*. 2021).

Seabirds from polar and subpolar regions are expected to be particularly sensitive to heat stress, because they have to maintain a thick body insulation in order to forage in cold waters (Davis & Darby 2012), but are increasingly exposed to high temperatures on land (Gimeno *et al*. 2025). It has been suggested that since marine birds have an “easy access” to water to cool down, they are unlikely to suffer much from hot conditions on land (Beaman *et al*. 2024). Yet, this assumption does not consider the high constraints that most seabirds face during reproduction, which is also the timeframe where the heat load is expected to be the highest. For instance, thermoregulatory behaviours such as bathing would directly impact nest attendance, thereby increasing the egg or chick exposure to heat and predators (Olin *et al*. 2023). Direct and indirect (*e.g.* longer foraging times, Bost *et al*. 2015) effects of heat may contribute to decrease reproductive success, leading to a trade-off between thermoregulation and reproduction (or even survival) in breeding adults.

The observation of thermoregulatory behaviours is an efficient and non-invasive way to study heat stress in free-living animals (Olin *et al*. 2023). Panting is considered a sign of heat stress as it is an energy-expensive evaporative cooling mechanism (Tan & Knight 2018). Panting behaviour during reproduction has been observed on days when air temperatures were high, for example in nesting guillemot (Olin *et al*. 2023), great skuas (Oswald *et al*. 2008) and African penguins (Welman & Pichegru 2022). The use of more passive thermoregulatory behaviours (*i.e.* less energetically costly than panting) has also been described in breeding seabirds, in order to increase convective and/or conductive heat loss for instance through spreading wings (Olin *et al*. 2023), bathing (Oswald *et al*. 2008), crouching or standing (Cook *et al*. 2020). To evaluate if thermoregulatory behaviours are sufficient to maintain normothermia, various methods exist for measuring body temperature in wild animals (for a review, see McCafferty *et al*. 2015). For animals being identified using sub-cutaneous pit-tags, the replacement of traditional pit-tag with thermosensitive ones offers the opportunity to collect subcutaneous temperature (T_sc_) while minimizing invasiveness (Andreasson *et al*. 2023; McCafferty *et al*. 2015). However, T_sc_ is measured close to the surface of the body and is known to be more variable than core body temperature (T_b_). Furthermore, the relationship between T_sc_ and T_b_ is also known to vary between species, and with the location of the sensor and environmental conditions (Lewden *et al*. 2017; Romanovsky 2018). Considering that measuring T_sc_ is usually less invasive and expensive than T_b_, it enables working with larger sample sizes, but requires prior validation in any given species and environmental context.

Here, we hypothesized that king penguins *(Aptenodytes patagonicus* J. F. Miller, 1778), the second largest penguin species, might be particularly sensitive to heat due to adaptation to cold waters, with effective body insulation for deep-diving (Davis & Darby 2012). Modifying this insulation during breeding is impossible, as penguins must alternate between incubating/brooding on land during spring/summer at fluctuating ambient temperatures (see ESM S1, numerous days with maximum temperature > 15°C at our study site), and foraging at sea in *ca.* 4°C water (Stonehouse 1960). The risks of heat stress are further accentuated as breeders reproduce on volcanic beaches with no access to shade, and must remain on land while fasting for prolonged periods, around 2-3 weeks during egg incubation and around 1 week during chick rearing (Weimerskirch *et al*. 1992). During their stay on land, king penguins breed in a dense colony, establishing a small territory against aggressive conspecifics (Côté 2000; Lemonnier *et al*. 2024) from which they cannot move or leave their egg/chick unattended without very high risks of reproductive failure due to predation (Descamps *et al*. 2005). This further accentuates the risk of heat stress by limiting access to cooling substrate (*i.e.* water) and triggering frequent antagonistic interactions (Viblanc *et al*. 2012). Although panting behaviour have been observed in polar and sub-polar penguin species (Stier & Lewden, *pers. obs.*), this phenomenon remains to be investigated in details.

In this study, we investigated the occurrence of heat stress in free-living king penguins and, by extension, assessed whether this species can serve as an early sentinel of the risk that sub-polar and polar seabirds face of experiencing land-based heat stress during reproduction in the context of climate change. To this end, we examined: (1) the occurrence of heat stress by studying thermoregulatory behaviours and the biological (*i.e.* breeding stage, sex and fasting duration) and climatic (*i.e.* ambient temperature, solar radiation, humidity and wind speed) factors triggering heat stress; (2) the relationship between subcutaneous temperature and core body temperature under increasing heat load (3) the relationship between thermoregulatory behaviours and subcutaneous temperature under hot climatic conditions; (4) the performance of different thermal indexes in predicting heat stress occurrence; and (5) the climatic conditions associated with reproductive failure.

## Materials and Methods

### Study model and site

We studied king penguins breeding in the “Baie du Marin” colony on Possession Island in the Crozet Archipelago (46°25′S; 51°52′E). Data were collected over two breeding seasons, 2021-2022 and 2022-2023. During breeding, male and female partners alternate between periods of foraging at sea with periods on land caring for the egg or chick (Figure S1). After laying a single egg, females return to forage at sea while males take care of the first incubation shift. Alternated guard continues throughout incubation (*ca.* 53 days) and brooding until the chicks reach emancipation (*ca.* 35 days) (Davis & Darby 2012). In this study, we monitored 72 breeding pairs (20 during 2021-2022 and 52 during 2022-2023). For each breeding pair, male and female parents were individually marked during courtship using hair-dye (Schwarzkopf Palette Deep Black) and monitored daily throughout incubation and brooding. The number of days on land (D_land_), from the return of the birds on land until the day of observation, was also considered in the analysis. The first incubation shift of males was excluded from the analysis as they fast 2-3 weeks longer than females, which influences greatly their physiology (Ancel *et al*. 2013).

### Climatic parameters

Climatic data were provided by Météo France through a weather station located *ca.* 1km from the penguin colony (Station 98404004, 46°25’S; 51°51’E). The station provided the air temperature under shelter (T_air_, °C), the hourly global solar radiation (SR, J/cm²), wind speed (WS, m/s), and relative humidity (RH, %). It also provided the dew-point temperature (Tw, °C). Although water vapor pressure is better than relative humidity to estimate the heat load and thermoregulatory constraints on an animal (Mitchell et al., 2024), it is highly colinear with temperature, and both cannot be included in the same model, contrary to RH. As our aim is to include the different meteorological variables within the same model, we thus used RH.

We used 4 different heat indexes to compare their predictive power to models including raw meteorological variables. These indexes are either practical metrics of dry heat load to which the biological responses of wild animals can be assessed (T_g_, Mitchell et al. 2024), the two most used indexes to predict heat stress occurrence in humans (THI, WBGT, Budd 2008), or two alternative indexes thought to better predict heat stress in humans and farm animals (HLI and WBGT; Carter *et al*. 2020; Liljegren *et al*. 2008; Silva *et al*. 2007). First, we estimated the globe temperature (T_g_) from climatic parameters provided by the weather station using an equation derived from field measurement of T_g_ in our penguin colony (T_g_, equation 1, see Figure S2 for details). The 3 other heat indexes we used were the Temperature-Humidity Index (THI, equation 2, Silva *et al*. 2007; Thom 1959), the Heat Load Index (HLI, equation 3, Gaughan 2002; Silva *et al*. 2007) and the Wet-Bulb Globe Temperature (WBGT, equation 4, Hajizadeh *et al*. 2017; Liljegren *et al*. 2008).

1. *Tg* = 2.89 + 0.83 × *Tair* − 0.035 × *WS* + 0.036 × *SR* (*see ESM S*2)
2. *THI* = *Tair* + 0.36 *Tw* + 41.5 *where T_w_ is the dew-point temperature*
3. *HLI* = 33.2 + 0.2 *RH* + 1.2 *Tg* − (0.82 *WS*) 0.1 − log(0.4 *WS*^2^ + 0.0001)
4. *WBGT* = 0.7 *Tnwb* × 0.2 *Tg* × 0.1 *Tair where T_nwb_ is the natural wet bulb temperature (for full equation, see Carter et al., 2020)*

### Thermoregulatory behaviours

Penguins’ thermoregulatory behaviours were recorded using scan sampling every 3 days, each scan encompassing 2 to 10 individuals and lasting 40 minutes, with an observation recorded every 2 minutes (21 observations per scan per individual). The scans were performed during midday (mean= 14:35 ± 1 hour and 35 minutes), when the heat load would be the highest, and to limit potential effects linked to circadian variation in behaviour. We gathered a total number of 1175 observations on N = 143 individuals, scattered over 60 days (from 28^th^ November 2021 to 8^th^ March 2022 and 4^th^ December 2022 to 26^th^ February 2023). The total number of scans per individual (mean of 9 ± 3 scans) varied with the duration of their shift on land (‘D_land_‘) and the survival of their offspring. We focused on 3 behaviours considered to have a thermoregulatory function, *i.e.* panting, wing spreading and exposure of the brood pouch (*i.e.* a highly vascularized and featherless area that may serve as a thermal window to dissipate heat; Handrich *et al*. 1997), and analysed their occurrence in terms of presence/absence during the scan (1/0 respectively). Observations were carried out by 11 observers during 2 field seasons.

### Body temperature measurements

We recorded sub-cutaneous temperature (T_sc_, °C) using thermosensitive transponders or “Pit-tags” (BioTherm13 tag, resolution ± 0.1°C, accuracy ± 0.5°C, between 33°C and 43°C, Biomark, USA) used for individual identification before the beginning of the study and placed in the upper back of all individuals (Oswald *et al*. 2018). T_sc_ was recorded every 3 days during shifts on land (n = 856), using the Biomark HPR Plus Reader just after behavioural scans and without handling of the animals. In our study species, T_sc_ changes have been shown to mirror changes in core body temperature while on land (Eichhorn *et al*. 2011; Lewden *et al*. 2017). Yet, to determine the extent to which T_sc_ can be informative of core body temperature (T_b_) in king penguins under high heat load, and thus of their actual thermoregulatory capacity, we used another set of data where T_sc_ was measured alongside T_b_ on 14 adults (7 males and 7 females, n = 37 observations) that are not part of the behavioural study. T_b_ was measured as stomach temperature using Anipill loggers (Bodycap®, Anipill, France, resolution ± 0.01°C, accuracy ± 0.2°C between 25 and 45°C) that were fed to the animals (Zuluaga *et al*. 2025). We estimated the raw correlation between T_b_ and T_sc_ recorded simultaneously, then calculated their difference (Diff = T_b_ - T_sc_, °C), and performed a Bland-Altman analysis (*Blandr* package, Deepankar Datta 2024), to assess the agreement between the two different measurements. To further investigate the implication of the different climatic variables on T_b_ and T_sc_ variations, we paralleled the data with the climatic variables corresponding to the same time and date (T_air_, SR, WS and RH). We tested the effect of climatic variables on T_sc_, T_b_ and on their difference using linear mixed models (with individual identity as random intercept).

### Reproductive failure

Survival of the egg or chick was monitored daily throughout the entire breeding season, and the day of egg abandonment or chick disappearance was recorded. The exact cause of reproductive failure (*e.g.* predation, parental abandonment) remained unknown in the vast majority of cases.

### Statistical analysis

Our behavioural observations and T_sc_ of breeding penguins were merged with climatic variables by the closest time of the day, with a maximum delay of 30 minutes. The statistical analyses were performed in R version 4.3.2 (R Core Team 2023). The effects of T_air_, SR, WS and RH on the probability of displaying the 3 thermoregulatory behaviours during the scans were analysed using generalised linear mixed models (R package ‘lme4’, version 1.1-35.1; Bates *et al*. 2015), assuming a binomial distribution (link=logit) and after scaling (*i.e.* z-transformation) of the climatic parameters. Linear mixed models with a normal distribution were used to assess the effects of climatic parameters on T_sc_. As individuals were measured several times over the breeding season, bird identity was included as a random intercept, as well as the identity of the observers (except for T_sc_, recorded by a pit-tag reader and therefore not subject to inter-observer variation). Year (2021-22 vs. 2022-23), sex (Male vs. Female), reproductive stage (Incubation vs. Brooding) and day of the shift on land (‘D_land’_) were included as fixed effects in the models. T_sc_ was further analysed depending on the presence or absence of panting behaviour for each individual during the preceding behavioural scan. Breakpoint analysis of T_sc_ against T_air_, SR, T_g_ and HLI, was performed using the *segmented* R package (version 2.0-2; Muggeo 2008). Segmented models were fitted from linear regressions without accounting for inter-individual variation of T_sc_, as not all individuals had enough data points to enable a mixed model breakpoint analysis. A Davies test was used to test for the significance of a change in slope in each linear model (Muggeo 2008).

Models including all variables but no interaction terms (*i.e.* No-interaction models) were compared to two-ways interaction models between climatic variables (*e.g.* T_air_*SR or WS*RH), and to a null model, using corrected Aikaike’s information criterion (AICc). The models with interaction terms presented in the results are the models retained based on the lower AICc. To assess model fit and statistical assumptions, ‘DHARMa’ package was used (version 0.4.6; (Hartig 2016). Variance inflation factors from the No-interaction models were all under 3, suggesting no substantial bias from collinearity between predictor variables (namely climatic variables) occurred (see Zuur *et al*. 2010). Conditional and marginal R² were used to compare the predictive power of each model (*i.e.* No-interaction model, Interaction model, heat indexes).

We tested for a difference in climatic conditions during days in which egg or chick mortality was observed, compared to days where no reproductive failure was recorded. For this, we used generalized models of the binomial family with the proportion of dead offspring each day depending on maximum T_air_, maximum SR, T_g_, and HLI. Maximum daily values were used since the exact time of day of reproductive failure was unknown, and maximum heat load was predicted to be more relevant than daily average. We also used non-parametric Wilcoxon tests to check for a difference in climatic condition between days with or without reproductive failure being observed.

## Results

### Behavioural thermoregulation

Independently of climatic conditions, panting was the most frequently observed thermoregulatory behaviour (23.4% of behavioural scans included panting behaviour), followed by wing spreading (9.7%) and exposure of the brood pouch (7.7%). Over the 60 days with behavioural scans, there were 27 days where at least one individual was observed panting (26 days for wing spreading and 16 days for brood pouch exposure).

The probability of observing panting behaviour was mainly explained by solar radiation (SR) and wind speed (WS), while air temperature (T_air_) and relative humidity (RH) did not significantly impact panting as main effects (Table 1A, Fig 1). Four two-ways interactions significantly explained panting probability, with the most notable being the interaction between T_air_ and SR, revealing that panting only increased with increasing T_air_ when SR were moderate to high (Fig 2A; see ESM S3 for details on other interactions). Males tended to display panting behaviour more frequently than females (only slightly, p = 0.067), and the probability to observe panting behaviour did not significantly vary with reproductive stage or between the two breeding seasons we studied (Table 1A). Panting probability significantly decreased with the time spent on land (D_land_), meaning that the birds’ susceptibility to suffer from heat stress gradually decreased after returning from the sea (Table 1A, ESM S4 A).

**Figure 1.**
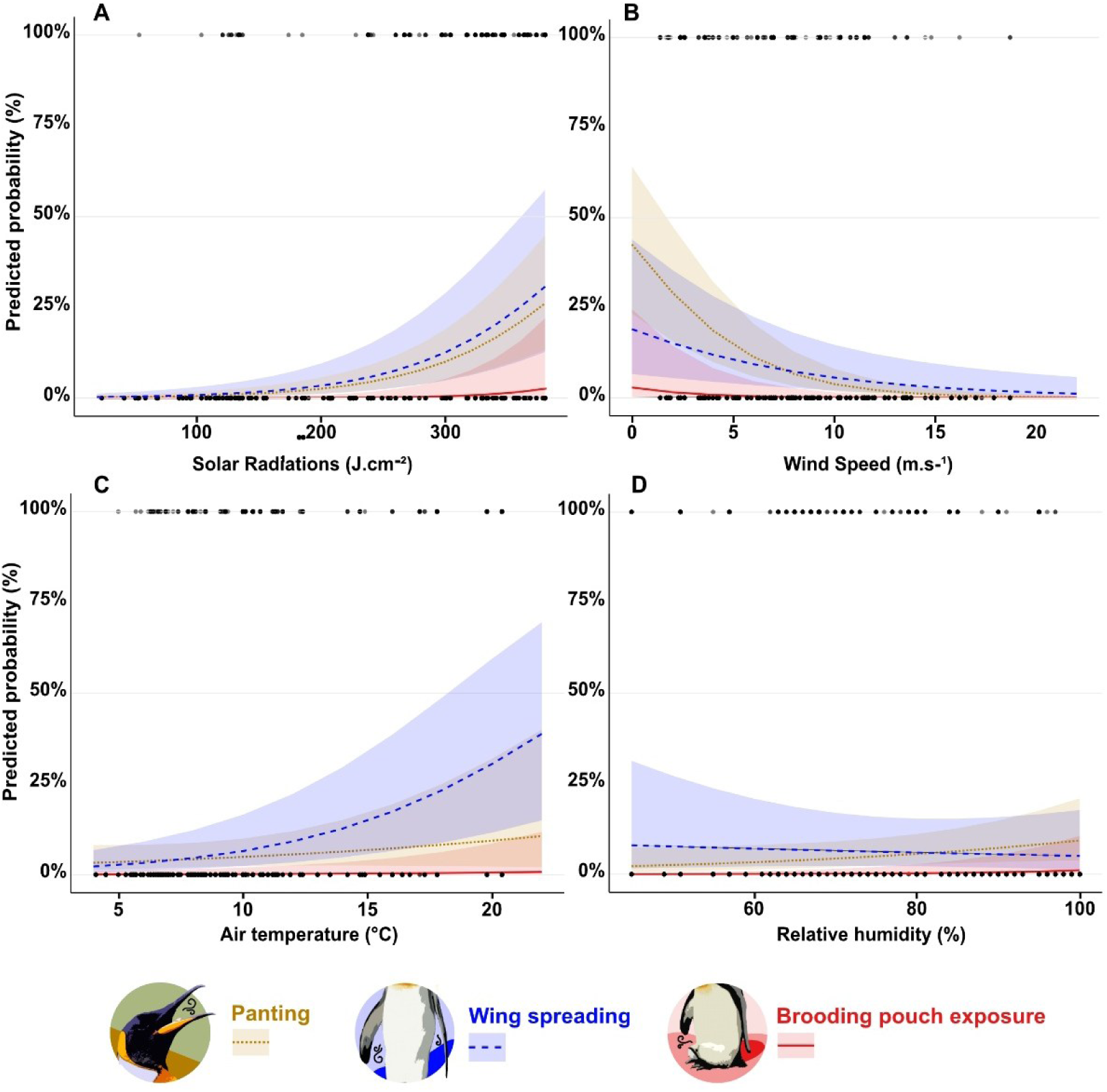
Predicted probabilities of thermoregulatory behaviours (panting (brown), wing spreading (blue) and exposure of the brood pouch (red)) occurrence according to (A) solar radiation, (B) wind speed, (C) air temperature and (D) relative humidity. Predicted probabilities were calculated from generalized linear mixed models; N = 144 individuals, n = 1175 observations.

**Figure 2.**
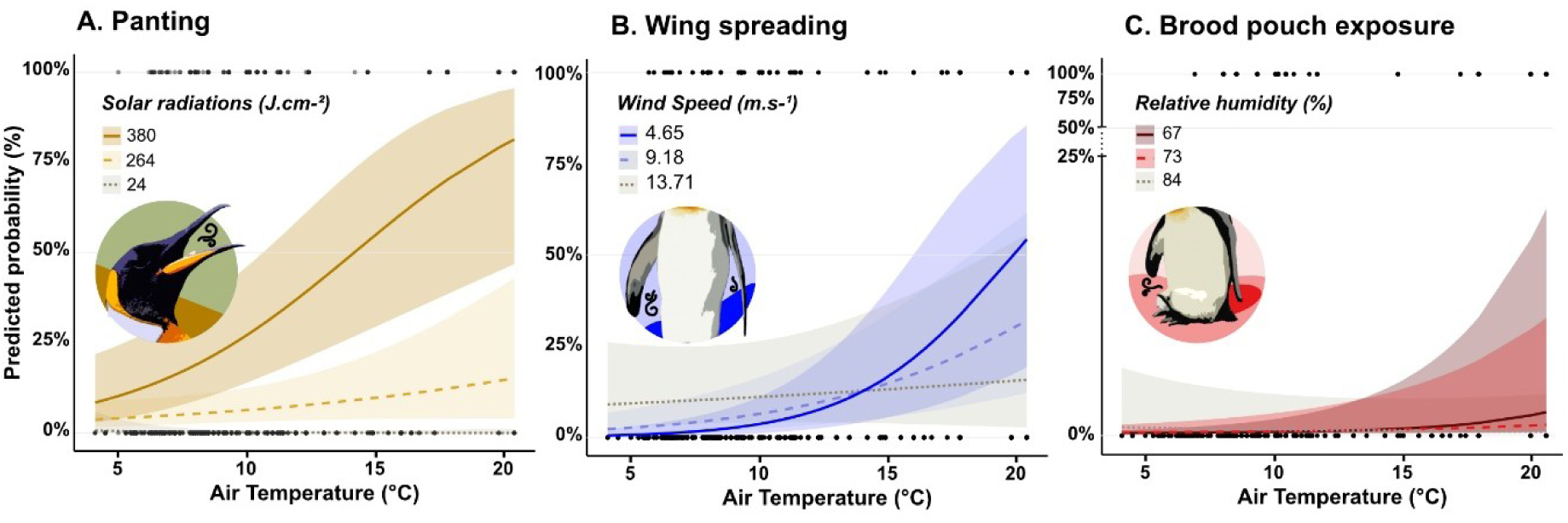
**Predicted probability of (A) Panting depending on air temperature and solar radiation;** solar radiation is represented as maximum (380), median (264) and minimum (24) registered during behavioural scans; (**B) Wing spreading according to air temperature and wind speed** (2^nd^ quartile (4.65), median (9.18), 3^rd^ quartile (13.71))**; (C) Brood pouch exposure according to air temperature and humidity** (2^nd^ quartile (67), median (73), 3^rd^ quartile (84)). Predicted probabilities were calculated from generalized linear mixed models; N = 143 individuals, n = 1123 observations.

**Table 1.**
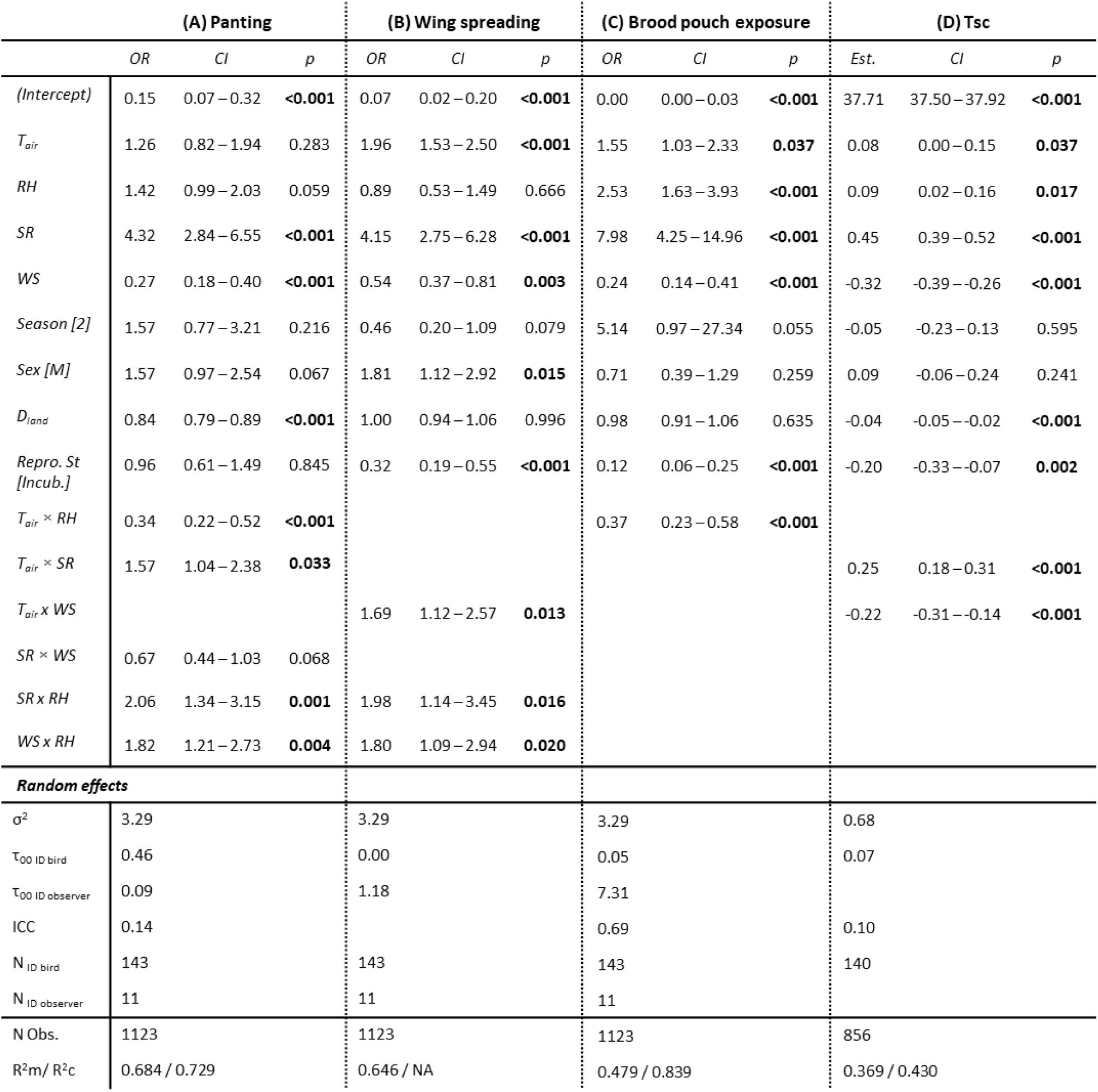
Summaries of the best fitted generalized linear mixed models predicting thermoregulatory behaviours: **(A) panting**, **(B) wing spreading**, (**C) exposure of the brood pouch,** and linear mixed model predicting **(D) sub-cutaneous temperatures** (Tsc) according to climatic parameters (air temperature: T_air_: solar radiation: SR; wind speed: WS; relative humidity: RH), their interactions, and biological parameters (sex [Male vs Female], season [2021-2022 vs. 2022-2023], reproductive stage [incubating vs. brooding], and days spent on land D_land_). For random effects, residual variance (σ²; intrinsically fixed to 3.29 for binomial models A, B and C), the between-subject variance (for penguin individuals Ʈ_00 ID bird_; for observers Ʈ _00 ID observer_), the global Intraclass Correlation Coefficient (ICC) and the number of individuals per random parameter are presented.

The probability of observing wing spreading increased under high solar radiation, high T_air_ and low wind speed, while relative humidity had no significant impact on this behaviour (Fig 1, Table 1B). Three two-ways interactions also significantly explained wing spreading probability, with the most notable being the interaction between T_air_ and WS (Table 1B), revealing that wing spreading increased with increasing temperature when wind speed was low to moderate (Fig 2B; see ESM S3 for details on other interactions). Males spread their wings significantly more often than females (p = 0.015) and brooding birds more than incubating ones (p < 0.001), but the probability to spread wings did not significantly change with the time spent on land nor between the two breeding seasons we studied (Table 1B).

The probability of observing the brood pouch lifted increased with SR, T_air_, RH and decreased with WS (Fig 1, Table 1C). The interaction between T_air_ and RH was also significant, revealing that brood pouch exposure only increased with increasing T_air_ when RH was low (Table 1C, Fig 2C). Sex, breeding season and the time spent on land did not significantly influence the probability of exposing the brood pouch, but brooding individuals were more likely to exhibit this behaviour than incubating ones (p < 0.001; Table 1C).

### Relationships between sub-cutaneous temperature, core body temperature and climatic conditions

In a subsample of king penguins (N = 14, n = 37) where both T_sc_ and T_b_ were simultaneously measured, T_sc_ was significantly correlated with T_b_ (r = 0.62, p <0.001, Figure S5A). The Bland–Altman analysis indicated that the two measures presented a nearly significant mean bias of 0.24°C (95% CI: –0.03 to 0.5, *t* = 1.81; *p* = 0.079), indicating that T_sc_ slightly underestimated T_b_, although not significantly (ESM S5B). However, the difference between T_b_ and T_sc_ significantly decreased with increasing body temperature (estimate of Difference ∼ Mean = −0.88, p <0.001; ESM S5B). The difference between T_b_ and T_sc_ significantly decreased with increasing T_air_ and SR (Table S6A and S6B, ESM S5C and D). Importantly, both T_sc_ and T_b_ showed the same pattern of increasing significantly with increasing T_air_ and SR (Table S6A and S6B, ESM S5C and S5D). Wind speed and relative humidity did not affect T_sc_, T_b_, nor their difference (Table S6C and S6D, ESM S5E and S5F).

### Relationships between subcutaneous temperature, heat load and thermoregulatory behaviour

Subcutaneous temperature (T_sc_) increased under high T_air_, SR and RH, and decreased in response to high WS (Table 1D). The best predictive model included significant interactions between T_air_ and SR and between T_air_ and WS (Figure 3, Table 1D), revealing that king penguin’s T_sc_ was especially elevated when T_air_ and SR increased concomitantly, along with a decrease in WS. Males and females did not show significant differences in their T_sc_, but brooding individuals had a higher T_sc_ than incubating ones (p = 0.002; Table 1D). T_sc_ also significantly decreased with days on land (p < 0.001, Table 1D, ESM S4B).

**Figure 3.**
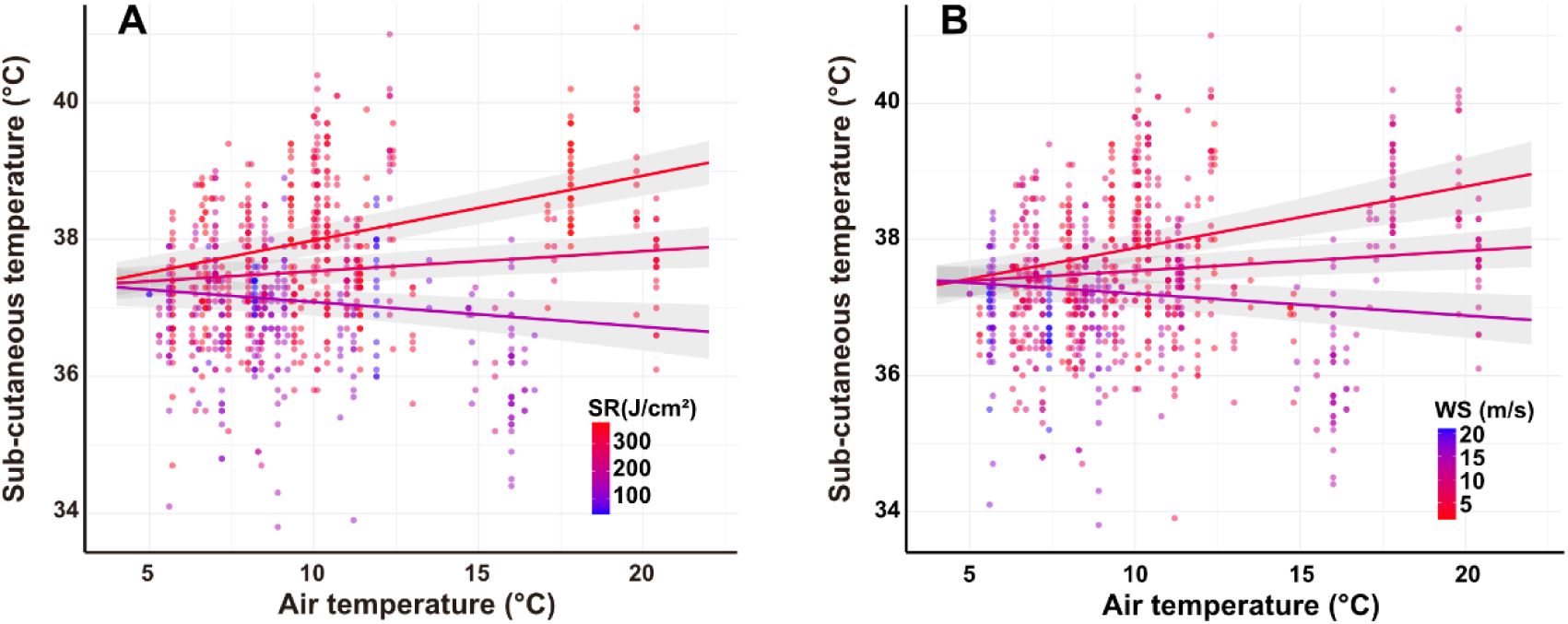
Breeding king penguins’ sub-cutaneous temperatures (T_sc_) according to (A) the interaction between air temperature (T_air_) and solar radiation (SR), and (B) air temperature and wind speed (WS). Values are predicted from the linear mixed model presented in Table 1D; *N = 140, n = 856*.

Breakpoint analyses showed that T_sc_ significantly increased with T_air_ when a threshold of 16.0 ± 1.1 °C was reached (Davies test: p = 0.005; ESM S7A), when SR reached 125.3 ± 11.7 J.cm-² (Davies test: p < 0.001; ESM S7B), or when the heat load index (HLI) was at 65.5 ± 1.4 (Davies test: p = 0.002; ESM S7D). However, the breakpoint of T_sc_ with increasing T_g_ was not significant (12.4 ± 0.6 °C; Davies test: p = 0.13; ESM S7C).

T_sc_ was significantly higher in panting than non-panting individuals (38.3°C vs. 37.2°C, t= 15.2, p < 0.001). Interaction models showed that T_sc_ increased with T_air_ and SR in panting individuals while this was not the case or less pronounced for non-panting animals (χ^2^ Panting*T_air_ = 7.5; p = 0.006; Figure 4A; χ^2^ Panting*SR = 14.3; p < 0.001; Figure 4B). Moreover, the increase of WS was not sufficient to decrease T_sc_ in panting individuals compared to non-panting birds (χ^2^ Panting*T_air_ = 10.2; p = 0.001; Figure 4C). Relative humidity had no effect on T_sc_ as a main effect, but the T_sc_ of panting individuals slightly decreased with increasing RH (χ^2^ Panting*RH = 4.3; p = 0.038 Figure 4D).

**Figure 4.**
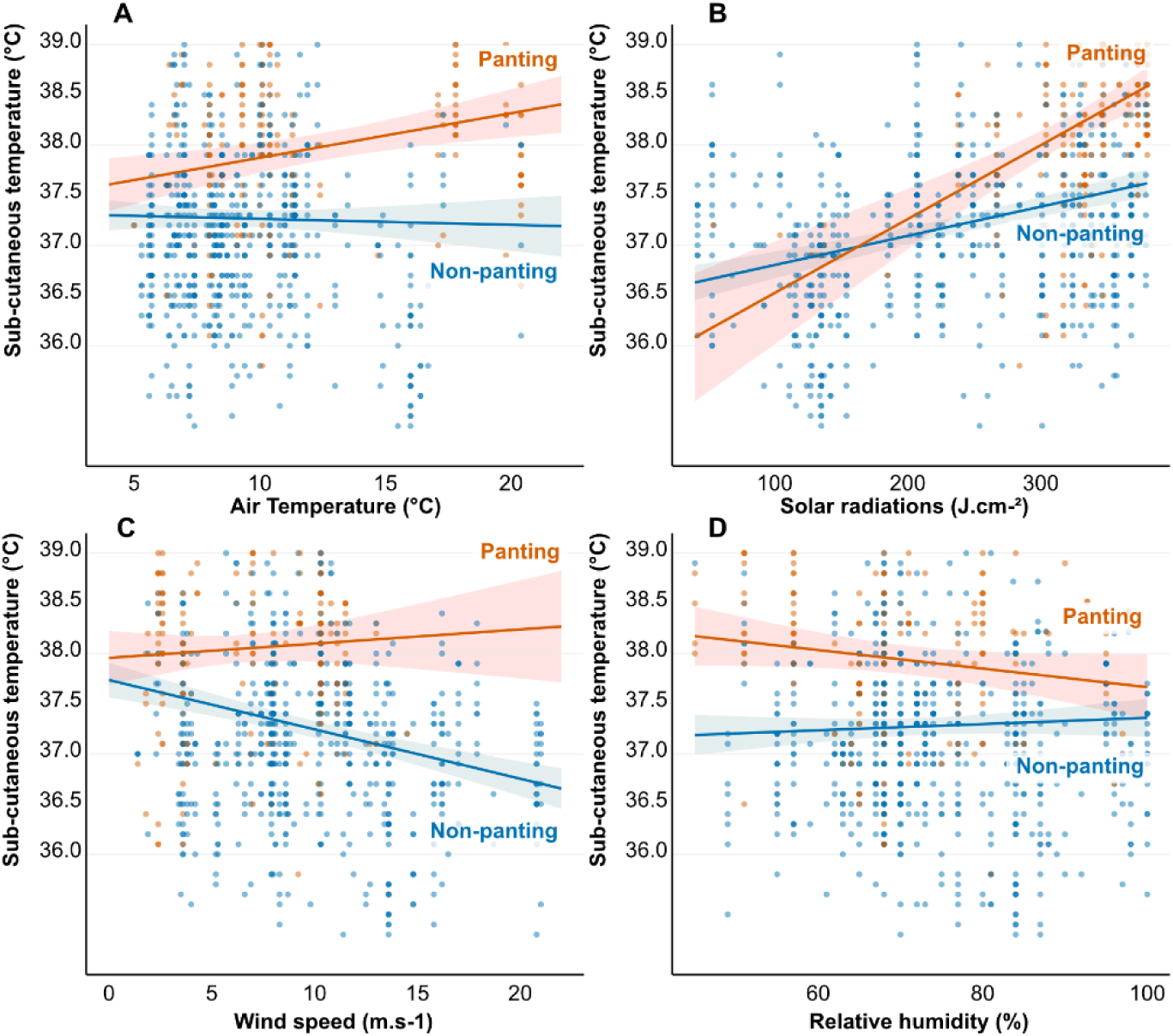
Relationship between air temperature (A), solar radiation (B), wind speed (C) or relative humidity (D) and king penguin’s sub-cutaneous temperature according to the use of panting as a thermoregulatory behaviour. *N = 140, n = 856*.

### Evaluating the relevance of T_air_ and heat indexes in predicting heat stress occurrence

Models only including T_air_ poorly explained the occurrence of thermoregulatory behaviours or variation in T_sc_, with R² ranging from 0.07 to 0.23 (Table S8). Solar radiation always performed better than T_air_ in predicting heat stress occurrence, with R^2^ > 0.24 (Table S8). Models containing all 4 parameters (T_air_, RH, SR and WS) as main effects (R² ranging from 0.30 to 0.67) and models including their two-ways interactions (R² ranging from 0.37 to 0.68) were the best predictive models for all explained variables (Table S8). Models including heat indexes (Globe temperature (T_g_), Temperature-Humidity (THI), Heat Load Index (HLI) and Wet-Bulb Globe Temperature (WBGT) explained less variance in thermoregulatory behaviours and T_sc_ than models (with or without interactions) including T_air_, RH, SR and WS as explanatory variables (Table S8). Among heat indexes, T_g_ (average R²m = 32.7%) and HLI (average R²m 32.0%) exhibited overall the best performance, while WBGT (average R²m 21.7%) exhibited intermediate and THI (average R²m 13.5%) the lowest performance (Table S8).

### Climatic conditions and reproductive failure

Over the two breeding seasons, 72 eggs were laid, 65 hatched and 53 chicks survived until emancipation, resulting in a 26.4% rate of reproductive failure (19 eggs or chicks that did not survive). Maximum T_air_ was significantly associated with a higher risk of reproductive failure (odds-ratio = 1.16; 95% CI = 1.02-1.32; p = 0.025; Table S9A), while maximum SR, Tg and HLI did not (Table S9B and D). The days where reproductive failure occurred (16 dates where parents abandoned, or the chick, sometimes several, were found dead) were significantly hotter than the days where no death was observed, but only regarding maximum T_air_ (T_air_ = 10.8 ± 3.3 °C, n = 170 for days without reproductive failure vs. 12.6 ± 3.4 °C, n = 16 for days with reproductive failure, W = 955.5, p = 0.025). Indeed, maximum SR did not differ between days with or without reproductive failure (SR = 283.8 ± 95.2 J.cm-² vs. 292.6 ± 73.6 J.cm-², W = 1582.5, p = 0.54), nor did maximum HLI (70.8 ± 4.6 vs. 71.8 ± 4.2, W = 1184, p = 0.20), or maximum T_g_ (T_g_ = 20.7 ± 4.3 °C vs. 21.9 ± 3.8 °C, W = 1183, p = 0.20).

## Discussion

Sub-Antarctic islands are expected to warm by 2.3 to 2.9°C over the next 100 years (Nel *et al*. 2023), and this increase in mean temperature does not necessarily reflect the positive trend of climate anomalies such as heat waves (Gimeno *et al*. 2025; Piatt *et al*. 2024; Stillman 2019), especially during the austral summer, *i.e.* the breeding season of many endemic polar and sub-polar species (Chambers *et al*. 2014). Our study unambiguously shows that king penguins do experience heat stress while breeding on land, at a frequency being far from anecdotal, with >20% of the behavioural scans at mid-day including signs of heat stress.

While many studies on heat stress focus on air temperature alone, the heat load experienced by wild animals markedly depends on other climatic variables (Mitchell et al. 2024). Here we show that the use of thermoregulatory behaviours in breeding king penguins was primarily influenced by solar radiation, followed by wind speed, air temperature and humidity in decreasing order of importance. Additionally, several interactions between climatic factors were significant in explaining the occurrence of heat stress, even if models without interactions were only slightly less predictive of heat stress occurrence than models including interactions (Table S8). Panting behaviour was the most frequently observed thermoregulatory behaviour, and occurred more often under high solar radiation and low wind speed, as well as at high air temperatures when solar radiation was moderate to high. Wing spreading was the second most frequently observed thermoregulatory behaviour, and it was more likely to occur under high solar radiation, high air temperature and low wind speed. Exposing the brood pouch was the least frequently observed behaviour, which can be explained by a trade-off between heat dissipation and the functions of the pouch: protecting the egg/chick from predators and keeping them warm. Yet, this behaviour was more frequent under high heat load (*i.e.* high solar radiation, high humidity, low wind speed and high air temperature), which supports our hypothesis regarding its potential use as a thermal window. High T_air_ has been reported to trigger heat stress in several seabirds and/or high latitude species (Cook *et al*. 2020; Holt & Boersma 2022; O’Connor *et al*. 2021; Olin *et al*. 2023; Welman & Pichegru 2022). While an increase in T_air_ as a main effect increased the probability to observe wing spreading or brood pouch exposure in breeding king penguins, it had no such effect on panting. Yet, air temperature in interaction with solar radiation explained panting behaviour frequency, to a point that virtually no bird would show panting around 20°C when solar radiation was low, while *ca.* 20% of the studied population was already observed panting below 10°C if sun exposure was maximal (Figure 3A). These results echo recent findings in guillemots, which showed an increased probability of panting with T_air_ in sun-exposed individuals around 15°C, compared with individuals in shaded areas in which panting started around 25°C (Olin *et al*. 2023).

Behavioural thermoregulation may enable animals to maintain normal body temperature under hot conditions, or only to limit hyperthermia (Talbot *et al*. 2017). Here, we first show in a sub-sample of 14 king penguins, that although sub-cutaneous temperature is more variable than core body temperature, they are moderately correlated, and both increase in response to high temperature and solar radiation. Yet, sub-cutaneous temperature over-estimates climatic effects on core body temperature, which is not surprising as peripheral tissues warm up at faster rates than core tissues due to heat gain from solar radiation and increased peripheral perfusion under high heat load to dissipate body heat through convection. While acknowledging these limits, subcutaneous temperature may still serve as a useful proxy of core body temperature when sampling a large number of individuals is required. Here, we showed that subcutaneous temperatures rose under increasing heat load (high solar radiation, air temperature and humidity, as well as low wind speed). Additionally, panting birds had a higher T_sc_ than non-panting ones, and T_sc_ increased with air temperature only in panting king penguins. This result may illustrate classic physiological thermoregulation: the panting birds that would be the most subjected to heat stress, underwent peripheral vasodilatation in order to evacuate body heat through convection, hence increasing T_sc_. The particularly thick feathered insulation coat of king penguins and limited thermal windows may reduce their capacity to lose heat through their body surface (Dawson *et al*. 1999; McCafferty *et al*. 2013). Additionally, breakpoint analysis allowed the detection of several thresholds above which T_sc_ would increase even further depending on T_air_, SR and HLI, which could illustrate a point at which core body temperatures increase. Although globe temperature has been suggested as an integrative metric of the environmental heat load (Mitchell et al., 2024), our analysis did not reveal a similar significant threshold for T_g_. Yet, to ensure the robustness of our conclusions, we would ideally need to measure core body temperature on a larger number of individuals to properly assess the penguins’ ability to maintain (or not) their core body temperature under high heat load. Our results contrast with studies in dovekies (*Alle alle*) suggesting that the core body temperature of this Arctic species is not challenged while attending the breeding colony on land (Grunst *et al*. 2023; maximum T_air_ = 13.5°C over 11 days) or in a captive set up to 25°C (*i.e.* substantially above the natural maximal temperature; Beaman *et al*. 2024). Yet, solar radiation was not investigated in the aforementioned studies, and empirical evidence, including here in king penguins, suggests that solar radiation is a key factor conditioning the total heat load faced by wild animals (Mitchell et al. 2024). Another factor that may contribute to explain these contrasted results between dovekies and king penguins lies in their difference in body size (ca. 0.15 vs. 12 kg), since smaller body size and thus larger surface area / volume ratio facilitates heat dissipation (Buckley 2021).

Taken together, the 4 climatic variables (T_air_, SR, WS and RH) we used predicted heat stress occurrence more accurately than T_air_ alone, or the four classical thermal indexes we used (T_g_, THI, HLI, WBGT). These indexes have been established for humans and farm animals (Gaughan 2002; Ioannou *et al*. 2023), which likely explain why they seem less suited for wild seabirds that present morphological and physiological adaptations to the cold, and breed in unsheltered conditions. Developing heat indexes specifically for wild endotherms would enable to better predict the response of natural populations to heat waves. Recent studies have predicted the increase in direct solar radiations (Ceppi *et al*. 2026), yet the local trends of cloud coverage and aerosols remain complex and depend on the regions and season (Goessling *et al*. 2025). As another example, near-surface wind speed in Antarctica has also been shown to either increase or decrease depending on the season and region (Andres-Martin *et al*. 2024). The complexity of meteorological trends, that depend on the geography, altitude, coastal or interior surfaces, or seasons, makes it difficult to forecast how species will effectively be challenged in the future during reproduction, especially since climate change predicts an increase in climatic anomalies (Gimeno *et al*. 2025). Considering the tremendous impact of solar radiation and wind speed on animal heat load, modelling future changes in climatic parameters beyond the sole air temperature seems of critical importance to predict the coping capacity of wild species in the context of climate change. This has for instance received recent attention in explaining water balance in wild zebra finches (Pacheco-Fuentes *et al*. 2025). Finally, for many species, general measures from meteorological stations may only reflect poorly the micro-climatic conditions experienced by animals, which depends on the habitat composition, and thus on-site measurement of micro-climate is needed (Mitchell *et al*. 2024), to better understand the importance of thermal refuges, involving factors such as soil conductivity, vegetation, shade or water (Gibson *et al*. 2026; Milling *et al*. 2018). Experimental studies manipulating for instance shading are, and will be, extremely useful in this context to better understand and quantify the impact of microclimate on animals’ response to hot environmental conditions (Corregidor-Castro *et al*. 2023).

Beyond climatic parameters, biological factors were also important in predicting the risks of king penguins to suffer from heat stress. First, sex did seem to play a role (albeit small) in explaining variation in heat stress occurrence, since males spread their wings more often and had a tendency to pant more than females. This may, at least partly, be linked to the higher size of beak and flipper relative to body mass found in females (Kriesell *et al*. 2018), which likely provides higher heat dissipation capacity through those thermal windows (Lewden *et al*. 2020). Second, brooding parents generally suffered more from hot conditions than incubating ones, as they spread their wings and exposed their brood pouch more often, as well as exhibited higher subcutaneous temperature than incubating individuals. Yet, brooding adults were not panting more than incubating ones. Considering that panting is extremely expensive in terms of water (Adélie penguins can lose water up to 0.3% of their body mass per hour while panting; Chappell & Souza 1988), our results may suggest that brooding individuals could limit the use of panting to avoid severe dehydration (Smit *et al*. 2016), especially because breeding king penguins do not appear to drink while fasting on land (Cherel *et al*. 1988). Brooding adults are also known to exhibit higher aggression rates towards conspecifics than incubating ones (Côté 2000), which could result in elevated body temperatures due to both physical activity and stress response (Knoch *et al*. 2022). Additionally, even if they are not fully thermally independent, growing chicks likely transfer some additional heat to the brooding adults (Duchamp *et al*. 2002). Finally, king penguins seemed especially sensitive to heat stress when freshly returned from the sea to incubate or brood on land. Indeed, they showed a higher probability to pant and a higher subcutaneous temperature during their first few days on land. This may be explained by the gradual transition to the hypometabolic state associated with fasting (2-5 days are needed to reach a stable hypometabolic state; Cherel *et al*. 1988), but also to their thermal acclimation to on-land conditions. The mechanisms involved in such a potential transient thermal acclimation on land remain to be investigated.

The impact of heat stress on breeding adult king penguins in our study was not lethal, contrary to what has been reported in Magellanic (with T_a_ reaching 44°C; Holt & Boersma 2022) and king penguins (two days at maximum T_a_ of 24°C and 30°C; Arriagada 2023) breeding in Argentina. Yet, sublethal effects on adult individuals may still have drastic consequences on population dynamics (Conradie *et al*. 2019), especially if reproduction is affected (Morosinotto *et al*. 2025), as we detected here. Reproductive failure may be related or accentuated by hot environmental conditions since parents may abandon reproduction because of heat stress, or be more susceptible to egg/chick predation when undergoing heat stress. Our results show that T_air_ was higher during days where reproductive failure occurred, which suggests that heat stress may be an important constraint on reproductive success for king penguins, as this has been demonstrated previously in common Guillemot (Olin *et al*. 2023). Yet, it was not possible in our study to determine whether the heat was directly (heat stroke) or indirectly (parent abandonment) lethal for the chicks, and it is surprising that only high T_air_ but not others parameters related to heat load was associated with reproductive failure. Better understanding the sublethal consequences of heat stress in king penguins, especially on reproduction, is thus needed to predict more accurately the changes in population dynamics expected due to the complex environmental challenges brought by climate change.

### Ethical note

All the procedures were approved by the French Ethical Committee (APAFIS#31268-2021042117037897 v3) and the Terres Australes et Antarctiques Françaises (Arrêtés TAAF A-2021-49 and A-2022-69).

## Data availability

The datasets and script are available on *FigShare*: https://doi.org/10.6084/m9.figshare.26968723

*A preprint is available on BioRxiv at* https://biorxiv.org/cgi/content/short/2024.09.09.611977v1.

## Author’s contribution

Study design: AS, VAV, AL. Funding acquisition: AS, JPR, PB, VAV. Data collection in the field: CL, CB, MM, MH, FB, EM, VAV, AS. Data analysis: AN. Writing original draft: AN and AS. Writing review and editing: AL, CL, CB, MM, MH, FB, EM, JPR, PB, VAV.

