## Supplementary materials for "The heat is on: behavioural, physiological and reproductive evidence of heat stress in breeding king penguins"

**Electronic supplementary material**

**ESM S1**

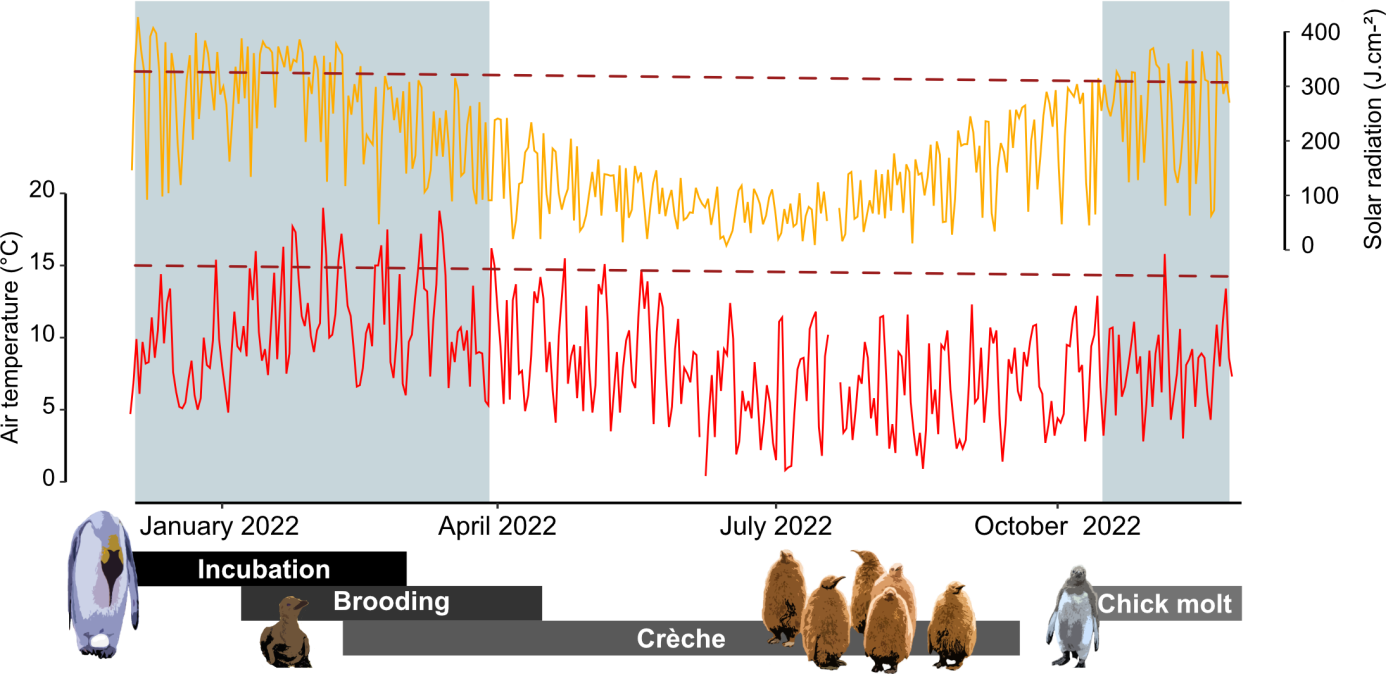

**Figure S1.** **Climatic conditions (daily maximum air temperature (in red) and maximum solar radiation (in orange) in 2022 in Crozet Archipelago along with a description of the breeding cycle of king penguins.**

**ESM S2. Predicting black globe temperature (*T*_g_) based on meteorological parameters**

Considering that T_g_ was not directly measured at our study site during the timeframe of this study, we relied on meteorological parameters to predict *T_g_*, based on actual *T_g_* measurements conducted during the 2023-24 field season with a Kestrel 5400 Heat Stress Tracker. We evaluated how measured T_g_ was best predicted by our four meteorological parameters (T_air_, SR, WS and RH) by selecting the best linear model using the R function dredge (Package MuMIn, version 1.48.11), which excluded RH from the model (adjusted R² of the selected model = 0.76).

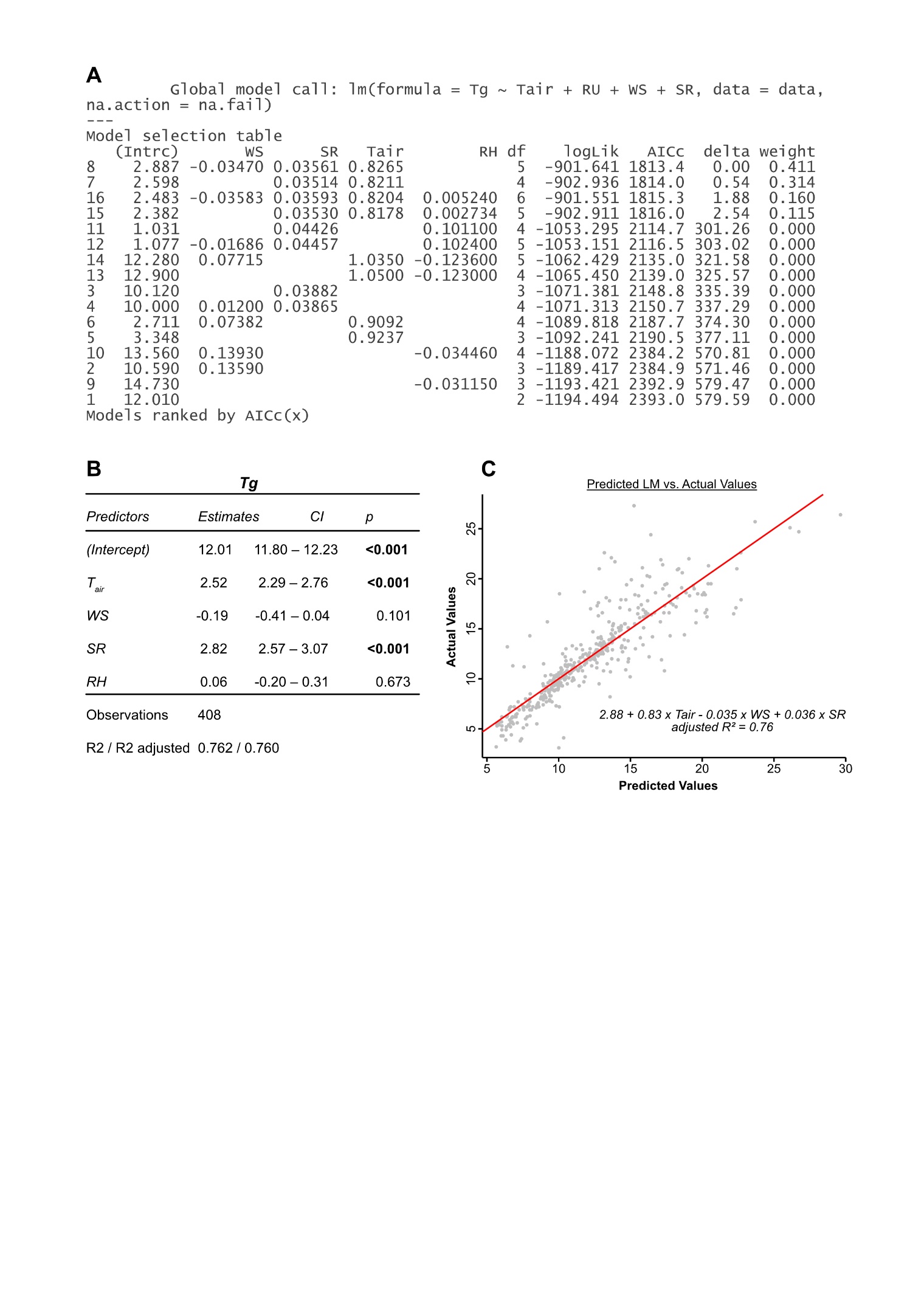

**Figure S2. Prediction of the globe temperature (T_g_) measured inside the penguin’s colony depending on meteorological parameters from the nearby Meteo France station (air temperature, solar radiation, wind speed and relative humidity), using linear modelization. (A)** dredge selection of lm(Tg~Tair+SR+WS+RH) depending on the lowest corrected Aikake’s Coefficient; **(B)** relative weight of the different climatic variables on globe temperature after scaling (z-transformation) and **(C)** Predicted vs. Actual values of T_g_.

**ESM S3**

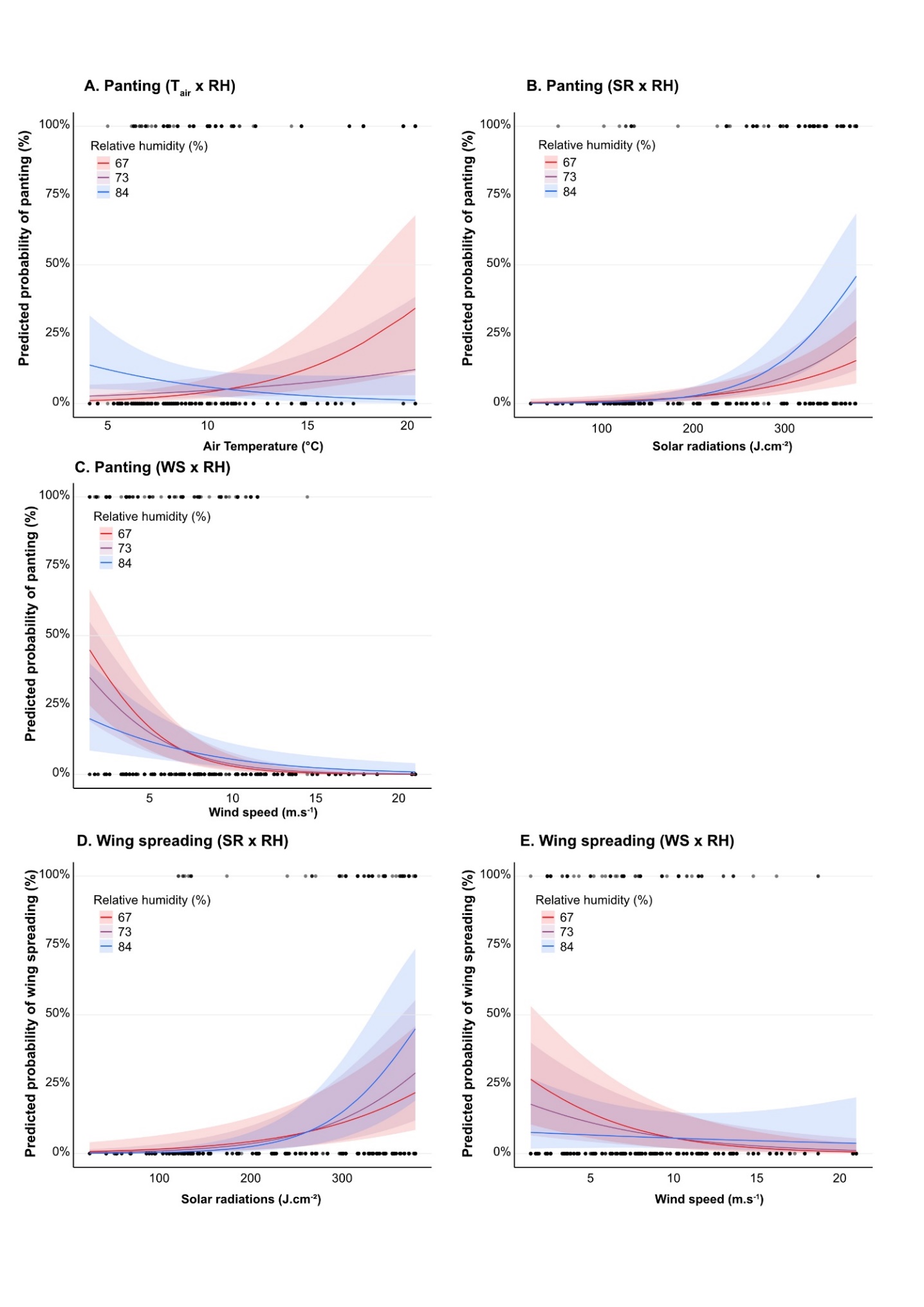

**Figure S3**. **Predicted probability of (A) Panting depending on air temperature and relative humidity** (**B) panting depending on solar radiation and relative humidity (C) panting depending on wind speed and relative humidity (D) wing spreading depending on solar radiation and relative humidity and (E) wing spreading depending on wind speed and relative humidity;** probabilities were calculated from generalized linear mixed models; N = 143, n = 1123 relative humidity (%) is represented as second quartile (67), median (73) and 3^rd^ quartile (84) registered during behavioural scans.

**ESM S4**

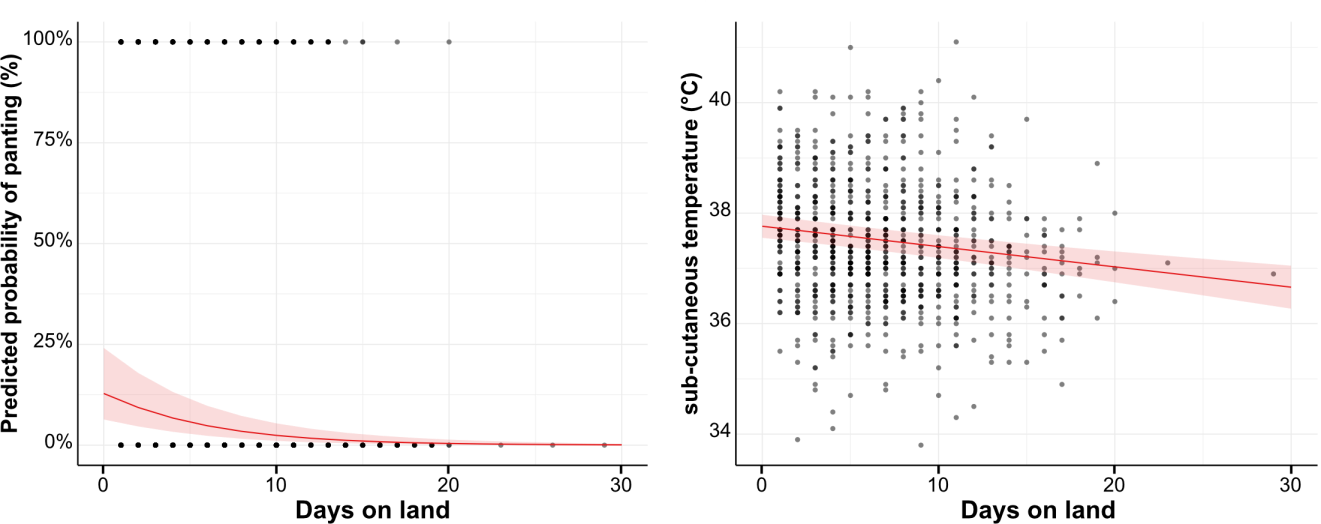

**Figure S4. Evolution of (A) panting probability and (B) subcutaneous temperature along the duration of the shift on land.** Values are predicted from linear mixed models presented in Table 1. N = 143, n = 1123 for panting; N = 140, n = 856 for T_sc_.

**ESM S5**

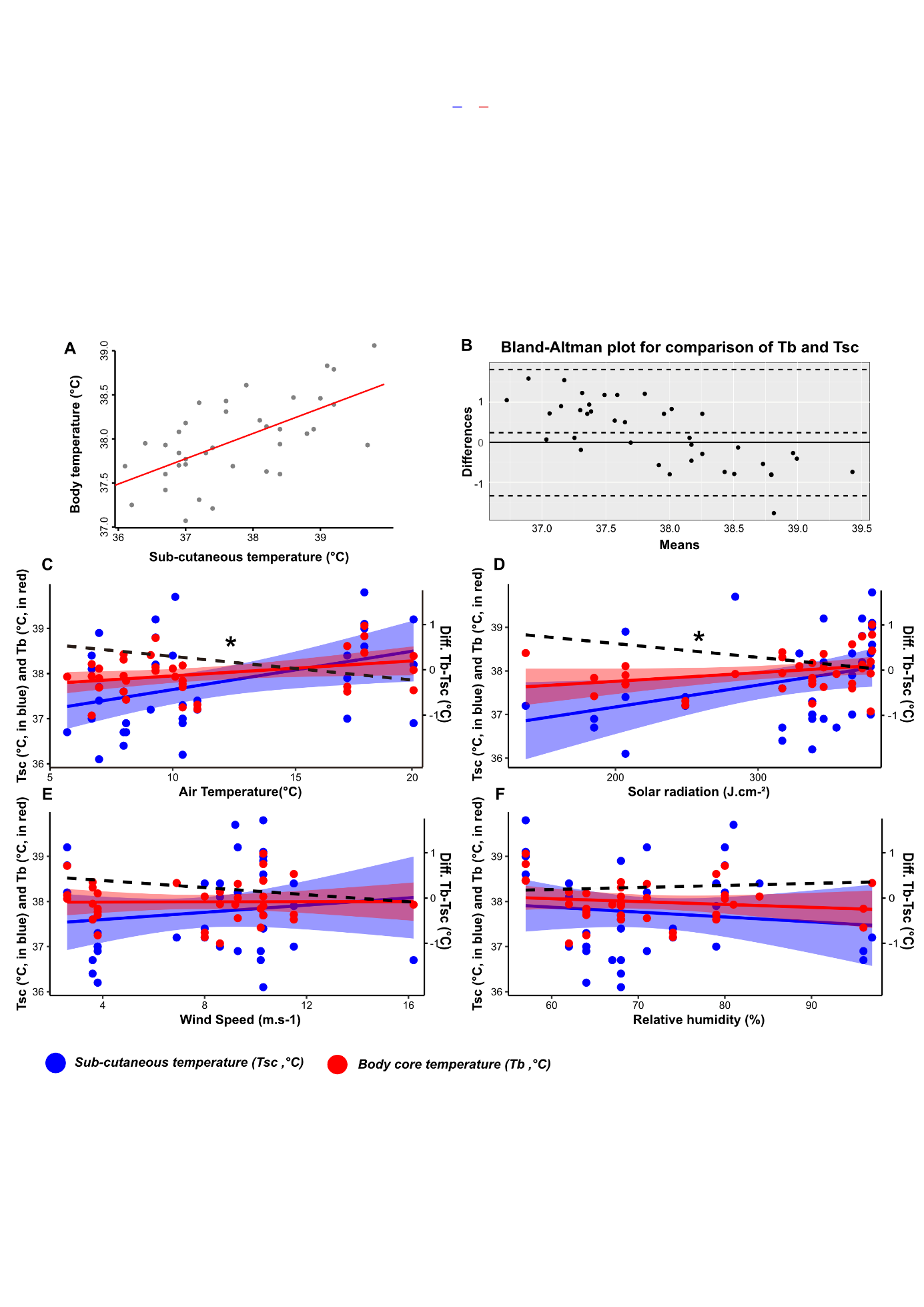
**Figure S5.** Relationships between **sub-cutaneous temperature (T_sc_, in °C) and core body temperature (T_b_, in °C). (A) Correlation (r = 0.62, p < 0.001), (B) Bland-Altman plot** representing T_b_ and T_sc_ difference (Ts_c_ – T_b_) of the same individuals against their mean, **(C-F) relationships between Tb (red), Tsc (blue) and their difference (T_b_ – T_sc_: black dotted line)** depending on climatic parameters (the significance of each relationship is marked with an asterisk): air temperature (°C), solar radiation (J.cm-1), wind speed (m.s-1) and relative humidity (%).

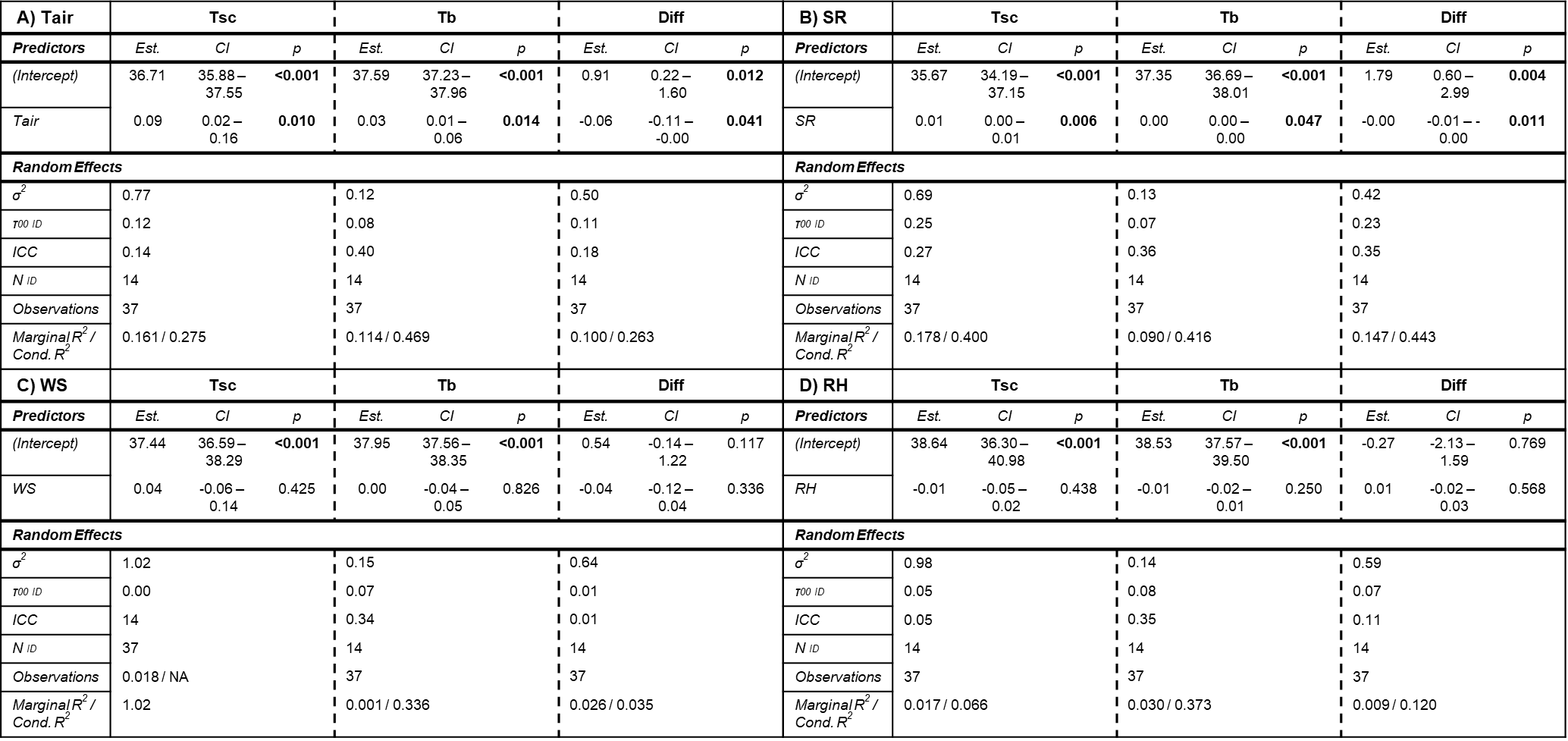
**ESM S6**

**Table S6. Summaries of the linear mixed models predicting sub-cutaneous temperature (T_sc_), core body temperature (T_b_) and their difference (Diff), depending on** **(A) ambient temperatures (T_air_)**, **(B) solar radiation (SR)**, (**C) wind speed (WS) and (D) relative humidity (RH**).

**ESM S7**

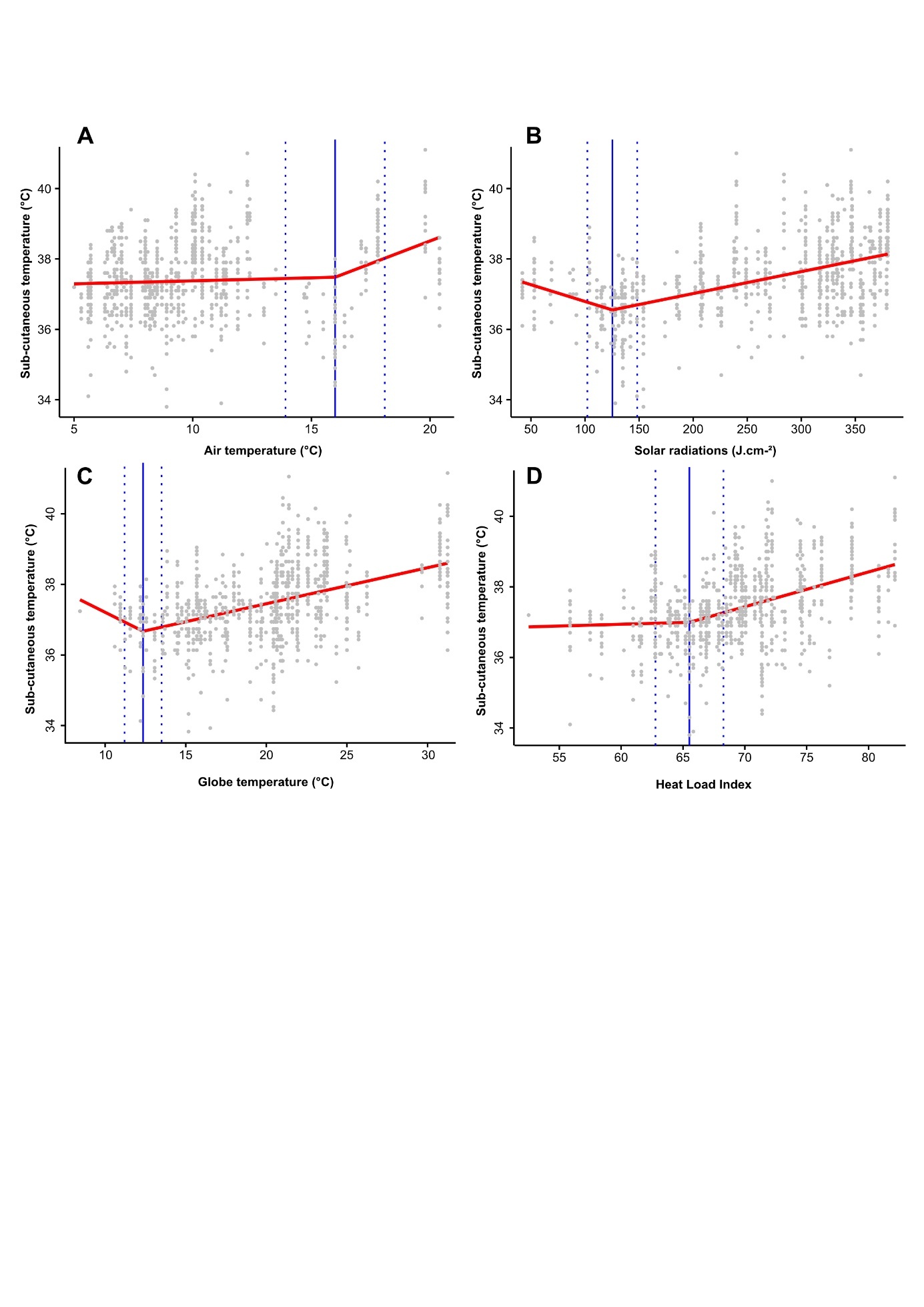

**Figure S7. Breakpoint analyses between king penguins’ sub-cutaneous temperatures (T_sc_) and (A) air temperature; (B) solar radiation; (C) Globe temperature and (D) heat load index (HLI)** computed from **air temperature, solar radiation, wind speed and relative humidity.** Segmented models are shown in red, and estimated breakpoints with confidence intervals are shown in blue. *N = 140, n = 856.*

**ESM S8**

**Table S8.** **Model comparison for the prediction of panting, wing spreading, brood pouch exposure and Tsc based on Akaike’s information criterion (AICc), marginal R² (R²m) and conditional R² (R²c).** The best predictive model with interactions (‘Inter.’) between all 4 climatic parameters (Tair, SR, WS and RH), is compared to a model without interactions (‘Total’), a model with only Tair or SR and 4 models including Heat indexes: globe

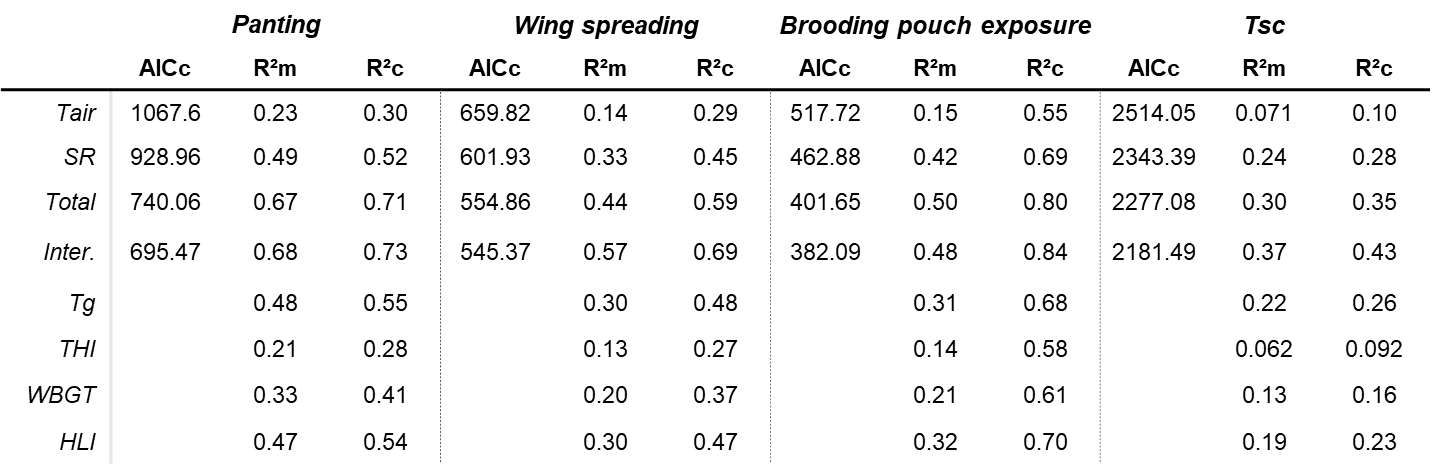

**ESM S9**

**Table S9.** **Summaries of the generalized linear models (binomial) predicting offspring death depending on**: **(A) maximum ambient temperatures (Tmax)**, **(B) maximum solar radiation (SRmax)**, (**C) maximum Globe temperature (Tgmax) and (D) maximum Heat Load Index (HLImax**).

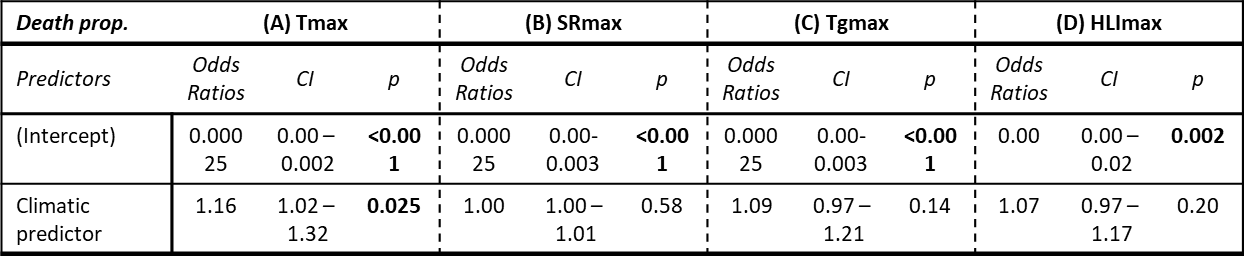
